# Treefrogs exploit temporal onset synchrony and harmonicity in forming auditory objects of vocal communication signals

**DOI:** 10.1101/2025.08.23.670182

**Authors:** Lata Kalra, Gerlinde Hobel, Mark Bee

## Abstract

Animals frequently communicate in dense social aggregations characterized by the presence of overlapping signals from multiple individuals. Receivers have to perceptually organize these overlapping signals into distinct ‘auditory objects’, each corresponding to the perceptual representation of an individual signal. All the signal components produced by the same individual should be integrated into a unitary auditory object while those produced by distinct individuals should be segregated. The principles of auditory object formation and their importance in vocal communication are less understood in non-human animals relative to humans. Here, using American green treefrogs, *Hyla cinerea,* we tested the hypothesis that receivers exploit the relative timing and harmonic relatedness of multiple spectral components to integrate or segregate sounds during auditory object formation. Using phonotaxis as a behavioral assay, females were given a choice between an attractive conspecific call consisting of three spectral components and a composite call consisting of the same three conspecific spectral components to which we added three spectral components from the call of a heterospecific sister species (*Hyla gratiosa*). The addition of heterospecific components to conspecific calls renders them less attractive. Across treatments, we manipulated the relative onset timing and harmonic relatedness of the heterospecific components in relation to the conspecific components. We predicted that asynchronous temporal onsets and inharmonic relationships between the conspecific and heterospecific components would promote their perceptual segregation. Our findings are consistent with this prediction. We discuss these findings in the light of parallel perceptual processes across animal taxa and neuroethological theories of auditory processing.

## Introduction

Animals often communicate in complex acoustic environments like noisy social aggregations, leks or choruses. Such environments are characterized by multiple signalers producing spectrally and temporally overlapping signals (Bee and Micheyl, 2008; Brumm and Slabbekoorn, 2005; Greenfield, 2005; McDermott, 2009). Behavioral decision-making requires a receiver to perceptually organize such overlapping signals into distinct ‘auditory objects’, corresponding to the perceptual representation of signals produced by distinct individuals (Bizley and Cohen, 2013; Bregman, 1990; Griffiths and Warren, 2004). Auditory object formation requires two complementary processes in which all the signal components (e.g., simultaneous harmonics and formants, or sequences of words, notes, and pulses) produced by the same signaler are perceptually integrated into a unified auditory object while those produced by different signalers are perceptually segregated into separate auditory objects.

Auditory object formation has been extensively studied in the context of speech recognition in noisy social environments. Various acoustic commonalities and differences between sounds can serve as ‘cues’ for auditory object formation in humans (reviewed in Bregman, 1990; McDermott and Oxenham, 2008; Moore and Gockel, 2012). Sounds tend to be integrated if they possess commonalities in their spectral and temporal features and if they originate from the same location. In contrast, sufficiently large differences in these same attributes promote segregation of sounds. Two such cues, temporal synchrony and harmonic relatedness (harmonicity) of sounds, frequently mediate perceptual integration of spectral components within speech signals (Darwin and Gardner, 1987). Speech signals constitute categories like vowels which, in turn, are composed of multiple formants (i.e., spectral regions with concentrated acoustic energy) that lend a vowel its distinct sound quality (Ladefoged and Disner, 2012). Formants with temporally synchronous onsets (ΔT = 0) or are harmonically related (ΔF = 0; are multiples of a common fundamental frequency) are likely to be perceptually integrated and heard as a single phonetic category. In contrast, sufficiently large deviations from these commonalities (ΔT ≠ 0; temporal asynchrony or ΔF ≠ 0; inharmonicity) promote segregation of formants from the vowel, thereby causing them to be heard separately from the rest of the vowel (Broadbent and Ladefoged, 1957; Brokx and Nooteboom, 1982; Darwin, 1981; Darwin, 1984; Darwin and Hukin, 1998; Gardner et al., 1989; Hukin and Darwin, 1995).

Non-human animals face the analogous challenge of communicating in complex acoustic environments using signals frequently composed of multiple concurrent spectral components (Fenton et al., 2014; Gerhardt and Huber, 2002; Hyland Bruno and Tchernichovski, 2019; Kershenbaum et al., 2016; Prestwich, 1994; Winn et al., 1981). Therefore, auditory object formation is crucial for accurate signal recognition, discrimination and localization in evolutionarily consequential contexts like survival and reproduction. Although auditory object formation is a ubiquitous communication challenge, our understanding of this process in non-human animal communication remains limited (Bee and Micheyl, 2008; Dent and Bee, 2018; Hulse, 2002). Non-human animals can employ similar auditory object formation cues as humans (Bee, 2010; Cai et al., 2018; Farris et al., 2005; Fay, 1998; Gupta and Bee, 2020; Klinge and Klump, 2009; Ma et al., 2010; Nityananda and Bee, 2011; Schul and Sheridan, 2006 *but also see* Bee and Riemersma, 2008; Kalra et al., 2024; Schwartz and Gerhardt, 1995; Schwartz and Serratto Del Monte, 2019). However, few animal studies have investigated multiple such cues in the same species. And as in humans (Elhilali et al., 2009; Micheyl et al., 2013b; Singh and Bregman, 1997), preliminary investigations of non-human animals suggest that multiple cues may be differentially weighed (Dent et al., 2016; Schwartz and Serratto Del Monte, 2019) or have additive effects (Itatani and Klump, 2020) on auditory object formation.

In this study, we investigated the role of temporal synchrony and harmonicity, two cues pivotal for speech perception in humans, in mediating auditory object formation in the American green treefrog (*Hyla cinerea*). Green treefrogs have been used extensively to investigate evolutionary (Ehret and Gerhardt, 1980; Gerhardt, 1987; Höbel and Gerhardt, 2003; Jones et al., 2014) and mechanistic (Fuzessery and Feng, 1983; Gerhardt et al., 1990; Lee et al., 2021; Schwartz and Gerhardt, 1989) aspects of vocal communication in complex acoustic environments. During their breeding season, male *H. cinerea* aggregate in dense choruses where they attract females by producing short (100-300 ms) advertisement calls that are repeated every 1-3 s during bouts of calling (Gerhardt, 1974a; Oldham and Gerhardt, 1975). Females select a mate by exhibiting phonotaxis toward a calling male and initiating physical contact that results in amplexus. The frequency spectrum of the advertisement call has a formant-like structure with acoustic energy concentrated in separate low-frequency (0.7–1.2 kHz) and high-frequency (2.4–3.6 kHz) bands. Most of the acoustic energy in the lower formant is concentrated in a single spectral component, whereas the higher formant typically emphasizes two to four spectral components (Gerhardt, 1974a,b; Gerhardt, 1976; Gerhardt, 2001; Oldham and Gerhardt, 1975). There is little acoustic energy between the two formants (1.2–2.4 kHz) (Fig. 1). Across much of its geographic range, *H. cinerea* frequently breeds in mixed-species choruses with the similar but somewhat larger barking treefrog (*Hyla gratiosa*). Green and barking treefrogs are closely related sister species that produce acoustically similar advertisement calls (Hua et al., 2009; Mecham, 1965; Robillard et al., 2006). The frequency spectrum of the *H. gratiosa* advertisement call also has a formant-like structure consisting of a lower formant and a higher formant, both of which are shifted to lower frequencies relative to the same formants in the *H. cinerea* call (Fig. 1). Females of *H. cinerea* do not avoid heterospecific (*H. gratiosa*) calls and will even approach them when given no other choice (Höbel, 2015). Importantly, however, they strongly prefer conspecific calls over heterospecific (*H. gratiosa*) calls (Oldham and Gerhardt, 1975). Moreover, the addition of spectral components from a heterospecific (*H. gratiosa*) call to a conspecific call starkly reduces the attractiveness of the signal relative to an unaltered conspecific call (Gerhardt, 1974b; Gerhardt and Höbel, 2005). The ability of simultaneous spectral components from heterospecific calls to render conspecific calls less attractive is key to the experimental design of the present study.

**Fig. 1.**
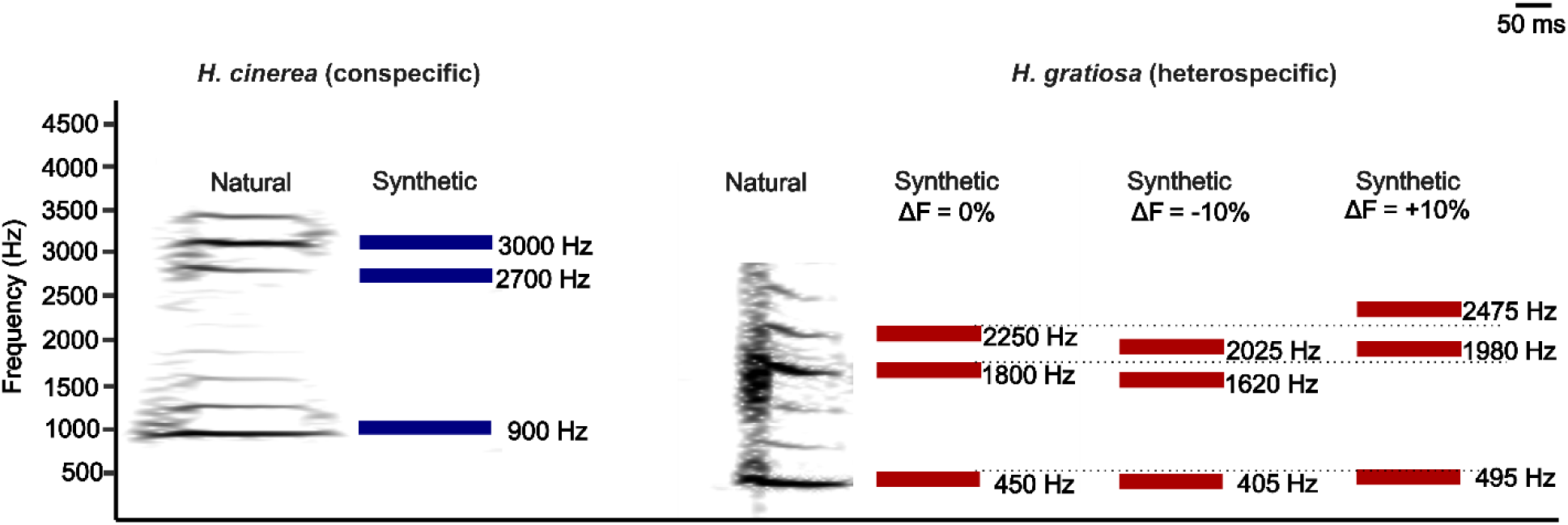
Natural and synthetic advertisement calls showing the formant-like spectra for *H. cinerea* (conspecific) and *H. gratiosa* (heterospecific). Sonograms depict the spectral components within a natural conspecific (left) and heterospecific (right) call. Here, greater intensity of the black shade depicts higher acoustic energy. Most of the acoustic energy is concentrated in regions around 900 Hz, 2700 Hz, and 3000 Hz in natural conspecific calls and around 450 Hz, 1800 Hz, and 2250 Hz in natural heterospecific calls. Based on the spectra of natural calls (Gerhardt, 1974a; Oldham and Gerhardt, 1975), synthetic advertisement calls were generated to have spectral components characteristic of *H. cinerea* calls (blue horizontal bars) and *H. gratiosa* calls (red horizontal bars). Three different sets of heterospecific spectral components were generated. One set (f_0_ = 450 Hz) was harmonically related to the conspecific spectral components (ΔF = 0%). The heterospecific components in the other two sets were mistuned, i.e., had their frequencies shifted either upward (495, 1980, and 2475 Hz) or downward (405, 1620, and 2025 Hz) by 10% (i.e., ΔF = ±10%) so that they were rendered inharmonic with the conspecific spectral components.

Using synthetic advertisement calls, we exploited the robust behavioral discrimination of female *H. cinerea* against heterospecific (*H. gratiosa*) spectral components to test the hypothesis that both temporal synchrony (ΔT = 0) and harmonicity (ΔF = 0) promote the perceptual integration of concurrent spectral components. In a series of two-alternative phonotaxis tests, we gave females a choice between a *standard* call with three simultaneous spectral components typical of conspecific calls (900, 2700, and 3000 Hz) versus a *composite* call consisting of the same three conspecific spectral components (900, 2700, and 3000 Hz) combined with three additional spectral components representative of the heterospecific calls of *H. gratiosa* (450, 1800, and 2250 Hz) (Fig. 1). The three conspecific components and the three heterospecific components each constituted a three-harmonic complex tone with fundamental frequencies (f_0_) of 300 Hz and 450 Hz, respectively. We capitalized on the coincidental property that, when combined, all six spectral components (450, 900, 1800, 2250, 2700, and 3000 Hz) were harmonics of a common fundamental frequency of 150 Hz. In a fully factorial design, we manipulated the degree of temporal synchrony and harmonicity between the conspecific and heterospecific spectral components of the composite call. These manipulations were done by shifting the onset timing and frequency of the heterospecific components relative to the conspecific components. We used the relative attractiveness of the standard and composite calls, as measured by selective phonotaxis, to assess the extent to which temporal synchrony and harmonicity promote perceptual integration (Fig. 2a). Based on earlier work in frogs (Gerhardt 1974; Gerhardt and Höbel, 2005), we predicted females would prefer the standard call under conditions (ΔT = 0 ms; ΔF = 0%) expected to promote integration of the conspecific and heterospecific spectral components of the composite call into a single (and less attractive) auditory object (Fig. 2a, b). Conversely, we predicted that introducing temporal onset asynchrony (ΔT ≠ 0 ms), inharmonicity (ΔF ≠ 0%), or both between the conspecific and heterospecific components of the composite call would promote their perceptual segregation; in turn, we expected perceptual segregation to render the composite and standard calls equally attractive based on their identical conspecific spectral components (Fig. 2a, c). Our factorial design allowed us to further explore potential interactions between temporal onset asynchrony and inharmonicity in promoting segregation.

**Fig. 2.**
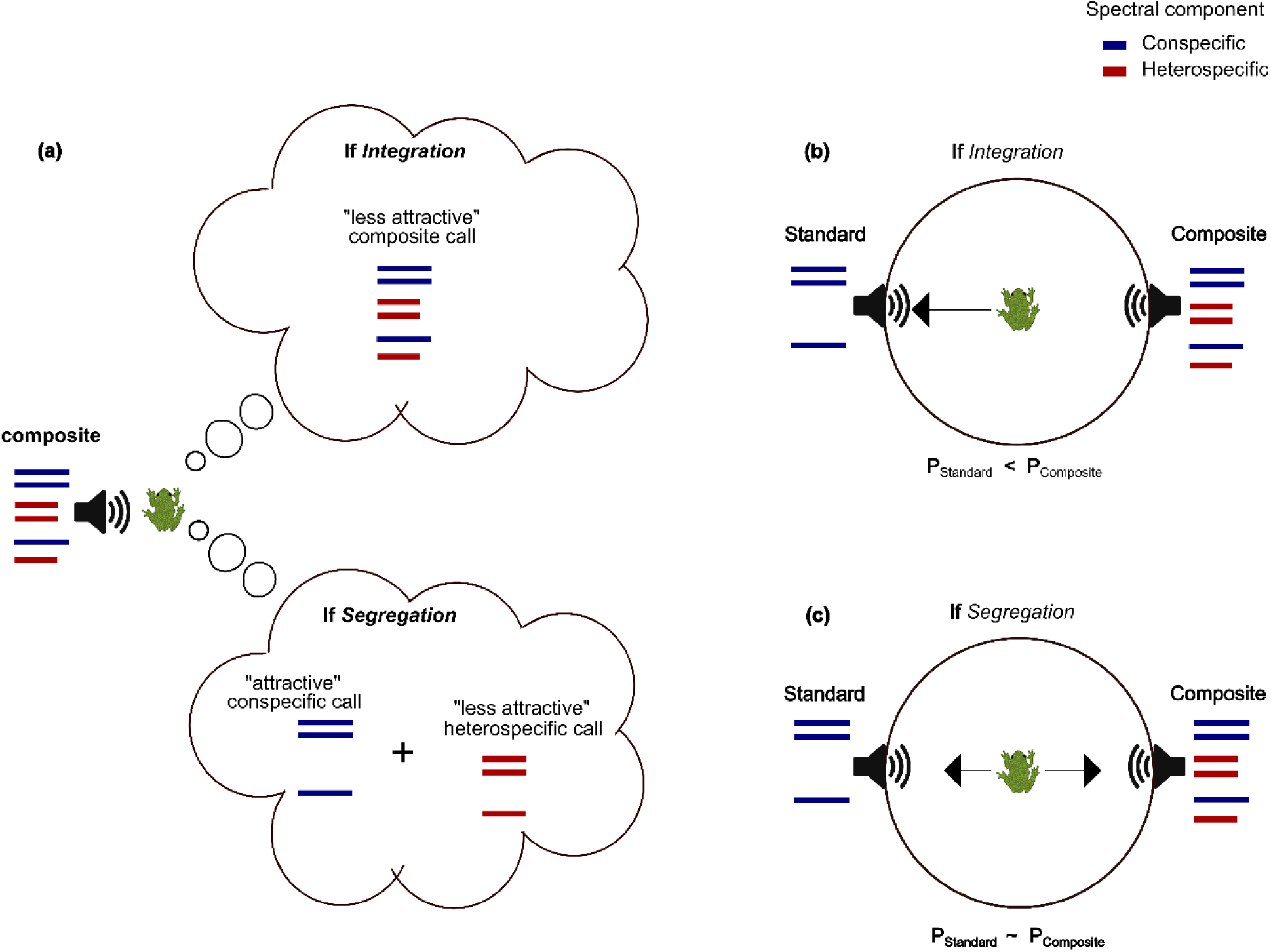
Integration versus segregation determines the relative behavioral attractiveness of the standard and composite calls. **a)** If subjects perceptually integrated the conspecific (blue bars) and heterospecific (red bars) spectral components in the composite call into a single auditory object, we predicted they would perceive the stimulus that to be less attractive than a standard call consisting only of the conspecific spectral components. In contrast, if subjects perceptually segregated the conspecific and heterospecific spectral components of the composite call into separate auditory objects, we predicted they would perceive two distinct calls, an attractive conspecific call (identical to the standard call), and a less attractive heterospecific call. Therefore, in a two-alternative choice test, we predicted that **b)** perceptual integration of the conspecific and heterospecific spectral components in the composite call would result in a greater preference for the standard call, whereas **c)** perceptual segregation of the conspecific and heterospecific components in the composite call would result in an equal preference for the standard and composite call based on both being perceived as having identical conspecific spectral components.

## Methods

### Subjects

Ninety gravid females of *H. cinerea* collected from Bowens Mill Fish Hatchery (Ben Hill County, GA, USA) were used as subjects in this study. Females were collected in amplexus at night (between 2100 and 2200 h) during June and July of 2019 and 2022. After collection, females were separated from their mates, transferred to separate containers, and subsequently tested in multiple phonotaxis tests (between 2200 h and 0500 h). There is little evidence of carryover effects when testing female frogs in multiple phonotaxis tests (Akre and Ryan, 2010; Gerhardt, 1981). Females and their mates were released at the collection site within 24 hours of collection.

### Stimulus and experimental design

Synthetic calls (44.1 kHz, 16 bit) were generated in MATLAB R2018b (Mathworks, Natick, MA, USA) and had spectral and temporal properties based on analyses of the natural advertisement calls of *H. cinerea* and *H. gratiosa* (Gerhardt, 1974a; Oldham and Gerhardt, 1975). We conducted a series of two-alternative phonotaxis tests in which females were allowed to choose between stimuli that acoustically simulated two males calling in perfect alternation, each at a rate of 40 calls/min. One stimulus consisted of repeated standard calls; the alternative stimulus consisted of repeated composite calls (Fig. 3). The standard call had a frequency spectrum that was typical of conspecific calls and is attractive to females (Gerhardt, 1974b). It comprised three spectral components (900, 2700, and 3000 Hz) that were 250 ms in duration (Fig. 3). The composite call combined the same three conspecific spectral components of the standard call with three heterospecific spectral components that were 150 ms in duration. The frequency of each heterospecific spectral component was determined by the level of ΔF (Fig. 3; Table 1). The shorter duration of the heterospecific components allowed us to manipulate, across treatments, the temporal onset asynchrony between the conspecific and heterospecific components while keeping the overall duration of the composite call consistent at 250 ms (Fig. 3; Table 1). The duration of the composite call was kept consistent across treatments to minimize any behavioral constraints imposed by subjects’ preferences for call duration *per se* (Gerhardt, 1987). The chosen durations of 250 ms and 150 ms for the conspecific and heterospecific components, respectively, fall within the natural range of call duration for both species (Gerhardt, 1974a; Oldham and Gerhardt, 1975). Note that, in designing the composite call, we encountered a trade-off between maintaining consistent call duration and consistent temporal ‘offset’ synchrony between the conspecific and the heterospecific components. As such, one limitation of this design was that, any shifts in the relative temporal onsets of the conspecific and heterospecific components introduced shifts in their relative temporal offsets (see below; Fig. 3). We chose to maintain a consistent duration of composite call over a consistent degree of temporal offset synchrony because temporal offset synchrony is believed to have a weak effect on perceptual organization (we elaborate on this argument further in the ‘Discussion’ section). Hereafter, nominal values of ΔT should be understood to correspond to the temporal onset asynchrony between the heterospecific components and the conspecific components.

**Fig. 3.**
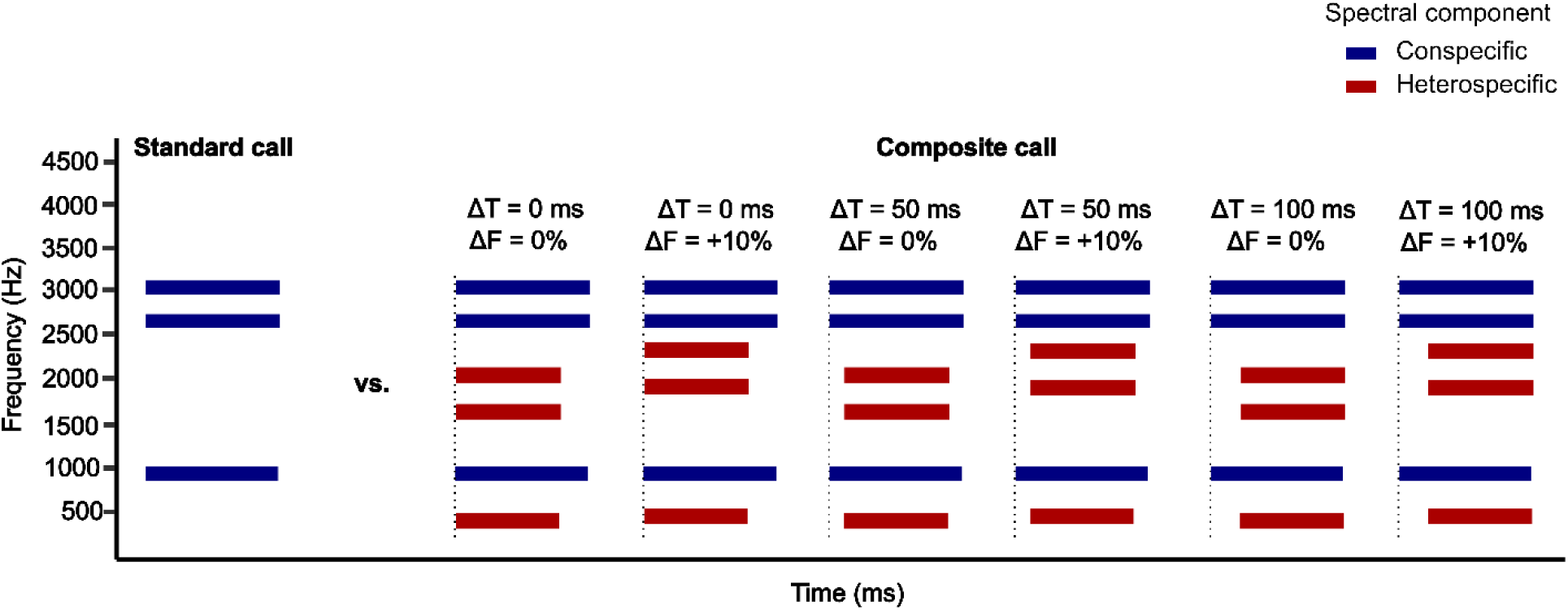
Exemplars of stimulus calls used across experimental treatments. The temporal onsets and frequencies of the heterospecific spectral components of the composite call were varied across treatments. Shown here are schematic exemplars of composite calls used in six of the 15 treatments depicting factorial combinations of three levels of ΔT (ΔT = 0, 50, and 100 ms; ΔT = 25 and 75 ms not depicted) and two levels of ΔF (ΔF = 0% and +10%; ΔF = -10% not depicted). Note that to manipulate ΔT (i.e., ΔT ≠ 0 ms), the onsets of the heterospecific components were delayed relative to those of the conspecific components. Consequently, there are corresponding shifts in the relative timing of the offsets of the conspecific and heterospecific components. In two-alternative choice tests, the standard call always alternated in time with one of the 15 possible exemplars of a composite call in a manner designed to simulate two antiphonally calling males.

**Table 1:** The 15 factorial combinations of ΔT and ΔF tested and the corresponding frequencies of conspecific and heterospecific spectral components.

| Treatment |  | Conspecific components | Heterospecific components |
| --- | --- | --- | --- |
| $\Delta F$ | $\Delta T$ | | |
| 0% | 0 ms | 900, 2700, and 3000 Hz | 450, 1800, and 2250 Hz |
|  | 25 ms |  |  |
|  | 50 ms |  |  |
|  | 75 ms |  |  |
|  | 100 ms |  |  |
| -10% | 0 ms | 900, 2700, and 3000 Hz | 405, 1620, and 2025 Hz |
|  | 25 ms |  |  |
|  | 50 ms |  |  |
|  | 75 ms |  |  |
|  | 100 ms |  |  |
| +10% | 0 ms | 900, 2700, and 3000 Hz | 495, 1980, and 2475 Hz |
|  | 25 ms |  |  |
|  | 50 ms |  |  |
|  | 75 ms |  |  |
|  | 100 ms |  |  |

In all stimulus calls, each spectral component had a random starting phase and an amplitude envelope shaped by inverse exponential rise and fall times of 25 ms and 50 ms, respectively. The starting phase of each spectral component in each stimulus call was re-randomized at the start of every night of testing, such that subjects tested on different nights heard different exemplars of the stimuli. At the beginning of each night of testing, each separate spectral component in each stimulus call was calibrated to a sound pressure level (SPL re 20 µPa; LCF) of 79 dB SPL at a distance of 1 m. Since a three-fold and six-fold increase in sound pressure corresponds to a 4.8 and 7.8 dB SPL increase, respectively (10 log(3) ∼ 4.8 and 10 log(6) ∼ 7.8), therefore, at 1 m, the overall level of the standard call (3 components) was 83.8 dB SPL and that of the composite call (6 components) was 86.8 dB SPL. These values fall within the natural range of variation in sound pressure levels recorded for both *H. cinerea* and *H. gratiosa* (Gerhardt 1975).

In total, there were 15 different choice tests based on the factorial combination of five levels of temporal asynchrony (ΔT = 0 ms, 25 ms, 50 ms, 75 ms, 100 ms) and three levels of inharmonicity (ΔF = 0%, +10%, -10%) (Table 1). We manipulated temporal asynchrony in the composite call by delaying the onset of its heterospecific components by ΔT relative to the onset of its conspecific components (Table 1; Fig. 3). Hence, at ΔT = 0 ms, all spectral six components had synchronous onsets and at ΔT = 100 ms they had asynchronous onsets and, consequently, synchronous offsets (Fig. 3). We manipulated harmonicity by either increasing (+) or decreasing (-) the frequency of each heterospecific spectral component by 10% (Fig. 1; Fig. 3). Thus, at ΔF = 0%, all six spectral components were harmonically related and shared a common f_0_ of 150 Hz. But at ΔF = ±10%, harmonicity within the three conspecific components and within the three heterospecific components was maintained, but the conspecific and heterospecific components were no longer harmonically related to each other (Table 1; Fig. 3). We tested ΔT within subjects and ΔF between subjects (n = 30 each for ΔF = 0%, +10%, and -10%; total n = 90), and we randomized the order in which different levels of ΔT were tested for each subject.

### Testing protocol

Phonotaxis tests were performed in a circular arena (2-m diameter and 60-cm height) that was located at the field site, approximately 300 m away from the nearest pond from which the subjects were collected. The arena’s floor was made of foam mats and its walls were constructed out of hardware cloth, covered in black fabric, to create a visually opaque but acoustically transparent barrier between the subjects and the speakers. Two speakers (Gallo Acoustics, Scotland, UK) were placed 180° apart from each other behind the arena wall, facing inward towards a subject’s release point at the center of the arena. Stimuli were broadcast using Adobe Audition 3.0 (Adobe Systems Inc. San Jose, CA, USA) installed on an HP ProBook 450 G6 (HP Inc., Palo Alto, CA, USA). Sound was output through a MOTU M4 sound card (MOTU, Inc., Cambridge, MA, USA), amplified by a Digital 2 Channel MOSFET amplifier (Dual Electronics, Corp., FL, USA), and broadcast through the two speakers. Sound pressure levels were calibrated using a Larson Davis Model 831 sound level meter (Larson Davis Inc., Depew, NY) connected to a microphone placed 1 m away from the speaker at the approximate position of a subject’s head at the release point. Across subjects, we randomized both the speakers assigned to broadcast the standard versus the composite calls as well as the type of call (standard versus composite) that was broadcast first.

At the beginning of a test, a single subject was placed inside a release cage fixed at the center of the arena floor and given an acclimation period of 60 s after which the stimulus broadcast began. During stimulus broadcast, the standard and composite calls alternated in time such that equal periods of silence preceded and followed each call. After two repetitions of both the standard and composite calls, the lid of the release cage was lifted remotely using a pulley system, and a timer was simultaneously started. After this point, the alternating calls continued while the subject was free to move inside the arena. A choice was recorded if the subject entered a 10-cm semicircular response zone in front of one of the speakers within 5 min. A no-response was recorded if the subject did not leave the release cage within 3 min or did not make a choice within 5 min. On average, females responded within 45.5 s of being released.

### Statistical analysis

Statistical analyses were performed in R studio version 2023.12.1 (α = 0.05 in all analyses). In our preliminary analysis, there were no significant differences in behavioral responses based on whether ΔF was positive or negative (see ‘Supplementary Materials’), we hereafter combine ΔF of +10% and -10% into a unified category of ΔF = ±10%. We used two-tailed binomial tests at each combination of ΔT (0 ms, 25 ms, 50 ms, 75 ms and 100 ms) and ΔF (0% and ±10%) to determine whether the proportion of subjects choosing the standard call exceeded the null expectation of 0.50. We assessed differential attractiveness of the standard and composite calls as functions of ΔT and ΔF by fitting a GEE model with logit-link function and an exchangeable correlation structure using the *geepack* package (Hardin and Hilbe, 2012). Wald-statistics were used to assess differential attractiveness as functions of ΔT and ΔF. Each subject’s choice was scored as a binary response variable (1 = standard call, 0 = composite call).

The independent variables were ΔT, ΔF, and their interaction. Since both ΔT and ΔF were coded as categorical variables, the output of the GEE model reported the effects of any combination of ΔT and ΔF in relation to a reference condition, which we specified as the test where ΔT= 0 ms and ΔF = 0%.

## Results

Subjects had a significant 4:1 preference (*p* < 0.01) for the standard call (proportion = 0.80) over the composite call (proportion = 0.20) when the conspecific and heterospecific spectral components of the latter had synchronous onsets (ΔT = 0 ms) and were harmonically related (ΔF = 0%) (Fig. 4, Table 2). In stark contrast, the proportions of subjects choosing the standard call were not significantly different from the null expectation of 0.50 when the conspecific and heterospecific spectral components of the composite call had asynchronous temporal onsets (ΔT ≠ 0 ms), were inharmonic (ΔF ≠ 0%), or both (Fig. 4, Table 2).

**Fig. 4.**
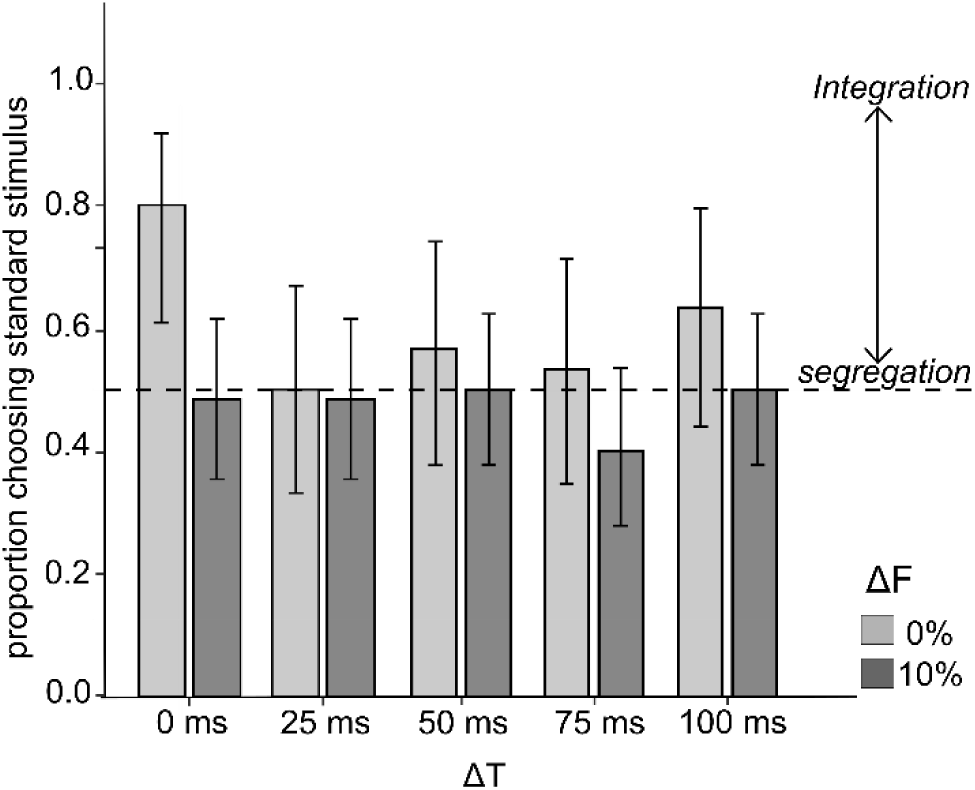
Proportion of subjects choosing the standard call across all combinations of temporal onset asynchrony (ΔT) and inharmonicity (ΔF). The dashed horizontal line indicates the null expectation of 0.50 in two-alternative choice tests. Gray and black bars indicate ΔF = 0% and ΔF = ±10%, respectively. Error bars represent 95% exact binomial confidence intervals. Temporal onset asynchrony (ΔT) was tested within subjects and inharmonicity (ΔF) was tested between subjects.

**Table 2:** Results of phonotaxis tests showing the proportions of subjects (±95% exact binomial confidence intervals) that chose the standard call over the composite call as a function of temporal onset asynchrony (ΔT) and inharmonicity (ΔF). Also shown are P values from two-tailed binomial tests of the hypothesis that the proportion choosing the standard call did not differ from the null expectation of 0.50.

| $\Delta T$ | $\Delta F$ | N | Proportion choosing the standard call | CI | P value |
| --- | --- | --- | --- | --- | --- |
| 0 ms | 0% | 30 | 0.80 | 0.61-0.92 | < 0.01* |
| | $\pm 10\%$ | 60 | 0.48 | 0.39-0.65 | 0.90 |
| 25 ms | 0% | 30 | 0.50 | 0.33-0.67 | 0.99 |
| | $\pm 10\%$ | 60 | 0.48 | 0.35-0.61 | 0.90 |
| 50 ms | 0% | 30 | 0.57 | 0.38-0.74 | 0.60 |
| | $\pm 10\%$ | 60 | 0.50 | 0.38-0.62 | 0.99 |
| 75 ms | 0% | 30 | 0.53 | 0.35-0.71 | 0.90 |
| | $\pm 10\%$ | 60 | 0.40 | 0.28-0.54 | 0.20 |
| 100 ms | 0% | 30 | 0.63 | 0.44-0.80 | 0.20 |
| | $\pm 10\%$ | 60 | 0.50 | 0.38-0.62 | 0.99 |

Compared to the treatment in which (ΔT = 0 ms, ΔF = 0%), introducing temporal onset asynchrony in the composite call while maintaining its harmonicity (i.e., ΔT ≠ 0 ms, ΔF = 0%) elicited a significant decrease in the proportion of subjects choosing the standard call for values of ΔT equal to 25 ms, 50 ms, and 75 ms, but not for ΔT equal to 100 ms. Additionally, introducing inharmonicity in the composite call while maintaining synchronous onsets (i.e., ΔF ≠ 0%, ΔT = 0 ms) also elicited a significant decrease in the proportion of subjects choosing the standard call compared to the reference test (Table 3). There was little evidence that introducing both temporal onset asynchrony and inharmonicity in the composite call resulted in strong interactions between these two perceptual segregation cues. In most factorial combinations where ΔT ≠ 0 ms and ΔF ≠ 0%, preferences for the standard call additively decreased (i.e., there was no significant interaction; Table 3). The only exception was the factorial combination of ΔT = 25 ms and ΔF = ±10%, for which there was a significant interaction because the proportion of subjects choosing the standard call was higher than that predicted from an additive decrease alone (Table 3).

**Table 3:** Results of the Generalized Estimating Equation (GEE) model. The effects of temporal onset asynchrony (ΔT), inharmonicity (ΔF), and their interaction are reported in relation to the reference test wherein ΔT = 0 ms and ΔF = 0%. Condition represents any manipulation in ΔT, ΔF or the interactive effects of different combinations of ΔT and ΔF. β indicates the estimate value for each condition. Wald Χ^2^ and P values correspond to the results of the Wald-statistics.

| Condition | $\beta$ | Wald $X^2$ | $P$ value |
| --- | --- | --- | --- |
| $\Delta T = 25$ ms | -1.39 | 5.35 | 0.02* |
| $\Delta T = 50$ ms | -1.11 | 4.37 | 0.04* |
| $\Delta T = 75$ ms | -1.25 | 5.08 | 0.02* |
| $\Delta T = 100$ ms | -0.83 | 1.95 | 0.16 |
| $\Delta F = \pm 10\%$ | -1.44 | 7.64 | 0.01* |
| $\Delta T = 25$ ms × $\Delta F = \pm 10\%$ | 1.39 | 3.90 | 0.05* |
| $\Delta T = 50$ ms × $\Delta F = \pm 10\%$ | 1.18 | 3.46 | 0.06 |
| $\Delta T = 75$ ms × $\Delta F = \pm 10\%$ | 0.91 | 1.97 | 0.16 |
| $\Delta T = 100$ ms × $\Delta F = \pm 10\%$ | 0.90 | 1.74 | 0.19 |

## Discussion

The primary goal of this study was to test the hypothesis that females of *Hyla cinerea* exploit temporal synchrony and harmonicity as perceptual cues in forming auditory objects of vocal communication signals. The data are consistent with this hypothesis. The main result from this study is that heterospecific spectral components rendered the composite call less attractive than the standard call *only* when they were temporally synchronous with (ΔT = 0 ms) and harmonically related to (ΔF = 0%) the conspecific spectral components. When the conspecific and heterospecific spectral components of the composite call had asynchronous onsets, inharmonic relationships, or both, the composite call was equally as attractive as the standard callwhich included only the conspecific spectral components. Our interpretation of these results is that temporal synchrony and harmonicity promoted the perceptual integration of the conspecific and heterospecific spectral components of the composite call into a single auditory object that was less attractive than a call consisting of only conspecific components. Onsets that were asynchronous by as little as 25 ms or shifts in frequency on the order of 10% promoted the perceptual segregation of the conspecific and heterospecific spectral components into different auditory objects. Because heterospecific calls are neither attractive nor inherently aversive in the presence of conspecific calls, onset asynchrony and inharmonicity reduced each female’s choice to one between two equivalent standard calls, for which there was no preference. Based on these results, we conclude that female green treefrogs exploit synchronous temporal onsets and harmonicity in forming auditory objects of vocal communication signals.

### Role of temporal onset synchrony in auditory object formation

Multiple studies in humans have demonstrated temporal synchrony as a robust cue for auditory object formation underlying the perception of speech perception as well as non-communication sounds. Bregman and Pinker (1978) and Dannenbring and Bregman (1978) presented human subjects with a sequence of a two-tone complex. When the temporal onsets of the two tones in the complex were synchronous, the tone complex was perceived as a single rich tone with a unified timbre. In contrast, when the temporal onset of the lower tone was shifted to lead the upper tone of the complex, the two tones were segregated and the distinct timbre was no longer perceived. Further investigations on vowel perception (Darwin, 1981; Darwin, 1984; Darwin and Ciocca, 1992; Darwin and Sutherland, 1984; Hukin and Darwin, 1995; Roberts and Moore, 1991) demonstrated that temporal onset synchrony effects the tendency of a single formant to be perceptually integrated with the rest of the vowel and, in turn, influences the perceived phonetic category of the vowel. For vowel sounds of shorter duration (∼ 50 ms) a temporal onset asynchrony as small as 30 ms was sufficient for perceptual segregation of a single formant from the rest of the vowel, while for longer vowel sounds, larger asynchrony was required. More recently, Elhilali et al. (2009) and Micheyl et al. (2013a) demonstrated that two tone sequences were perceptually integrated and heard as a ‘single’ sequence when the tones shared temporal synchrony. Shifting the temporal synchrony between those tones promoted the segregation of sequences thereby causing the percept of two distinct sequences.

Thus far, only two studies have investigated if non-human animals can exploit temporal synchrony in forming auditory objects of communication signals. Neilans and Dent (2015) presented budgerigars with synthetic conspecific contact calls and demonstrated that increased temporal asynchrony between overlapping conspecific signals promoted their perceptual segregation, thereby causing them to be heard as separate signals. Gupta and Bee (2020) demonstrated, in Cope’s gray treefrogs, that the two spectral components of conspecific calls were integrated when they were temporally synchronous. Our study corroborates both of these findings and thus contributes to the emerging evidence that temporal synchrony may be a robust auditory object formation cue in non-human animals. Additional studies are needed to assess the role of temporal synchrony in perceptual organization in other non-human animal species.

It is also worth looking at the present findings in the light of male-male signaling dynamics in nature. Males in dense acoustic aggregations frequently adjust the relative timing of their signals in two broad ways. First, commonly seen in birds, frogs and insects, the relative signal timings are asynchronous so as to approximate alternation or minimal signal overlap between the signals of different signalers (reviewed Gerhardt and Huber, 2002; Hall, 2009; Klump and Gerhardt, 1992; Thorpe, 1975). Second, a relatively uncommon signaling pattern is one in which the relative signal timings are synchronized so that there is maximal signal overlap between the signals of different individuals (Grafe, 1999; Legett et al., 2020; Legett et al., 2021; Tuttle and Ryan, 1982; Wells and Schwartz, 1984). Signal alternation might serve to facilitate a signaler’s assessment of a neighbor’s call, minimize degradation of species-typical temporal information required for mate choice or exploit a female’s perceptual bias to prefer a leading male’s calls (Greenfield et al., 1997; Schwartz, 1987). On the other hand, signal synchronization may serve to attract females from a distance, avoid predators by potentially impairing their ability to localize individual signalers, or to jam the signals of neighboring males (Grafe, 1999; Greenfield, 1994; Greenfield et al., 1997; Legett et al., 2021; Tuttle and Ryan, 1982). There is emerging evidence that adaptive significance of different signaling dynamics may relate to the perceptual processes within the receivers (Bosch and Márquez, 2002; Grafe, 1999; Höbel and Gerhardt, 2007; Minckley and Greenfield, 1995; Reichert et al., 2024; Snedden and Greenfield, 1998). Neilans and Dent (2015) and the present study demonstrated that temporal asynchrony between signal produced by different individuals promotes the perceptual segregation of these signals by the receiver, making the signals easier to recognize. Given the diversity of inter-male calling dynamics across animal taxa, more work would be needed to investigate how the temporal patterns of such signaling interactions relate to perceptual organization processes in conspecific and heterospecific receivers.

One limitation of this study was that it did not control for the possible influence of temporal offset asynchrony. Because female green treefrogs have duration-dependent preferences, we elected to maintain a consistent duration of the composite call across treatments versus a consistent temporal offset synchrony of the conspecific and heterospecific components within the composite call. Because the conspecific and heterospecific components had unequal durations, any change in their relative temporal onsets was necessarily coupled with a change in their relative temporal offsets. For instance, at a temporal ‘onset’ asynchrony of 0 ms, the conspecific and heterospecific spectral components had a temporal ‘offset’ asynchrony of 100 ms. We speculate that any behavioral effects imposed by the uncontrolled temporal offsets in this study were marginal for two reasons. First, at least in humans, asynchronous temporal onsets have a significantly greater effect on promoting perceptual segregation than asynchronous temporal offsets (e.g., Darwin 1984; Madsen and Moore 2014). Second, our results indirectly suggest that perceptual segregation was primarily driven by asynchrony in temporal onsets. While the proportions of subjects choosing the standard call did not differ significantly depending on whether spectral components in the composite call had asynchronous onsets (ΔT ≠ 0 ms) versus asynchronous offsets (ΔT ≠ 100 ms; see Table 3), only asynchronous onsets (ΔT ≠ 0 ms) yielded significant preferences for the standard over the composite call (Table 2). Follow-up studies could adopt an alternate design in which the offsets of the conspecific and heterospecific spectral components are kept synchronous and onset asynchrony is manipulated by varying the duration of the heterospecific spectral components.

### Role of Harmonicity in auditory object formation

Research using synthetic tone-complexes and speech sounds reveals that harmonicity mediates perceptual organization of concurrent spectral components. Darwin (1981), Darwin and Gardner (1986) and Gardner et al. (1989) demonstrated that mistuning a single formant within a vowel causes the formant to be perceptually segregated from the rest of the vowel, leading to an overall change in the perceived timbre of the vowel and, in turn, the perception of the phonetic category of the vowel. Additionally, Moore et al. (1985, 1986) and Hartmann et al. (1990) demonstrated that mistuning single components within a harmonic complex causes them to be perceptually segregated and heard separately from the tone-complex. A mistuning as small as 3% could be detected by subjects in the form of perceived changes in the amplitude envelop of the tone complex, whereas mistuning by greater than 8% led to complete perceptual segregation.

Even though signals in non-human animals are frequently characterized by harmonic complexes that resemble vowel sounds (Hoeschele, 2017; Kershenbaum et al., 2016), few studies have investigated how harmonicity impacts perceptual organization of communication signals. Lohr and Dooling (1998) used stimuli modelled on natural calls and showed that both zebra finches and budgerigars are highly sensitivity to the mistuning of single harmonics within calls. Simmons and Bean (2000) found that the evoked calling response of male bullfrogs was significantly lower when particular harmonics within the call were mistuned, compared to when all the frequencies within the signal were harmonic. Klinge and Klump (2009, 2010) demonstrated that gerbils were remarkably sensitive to mistuning of individual components within a tonal complex. In those studies, shifts in behavioral responses were observed even when inharmonicity was introduced by very small shifts (< 1 Hz) in the frequency of individual components. Our findings, demonstrating that perceptual segregation is promoted upon mistuning of the specific spectral components corroborates these previous findings on birds, frogs and rodents and adds to the evidence for the prevalence of harmonicity mediated perceptual organization in non-human animals. In conjunction with our findings on temporal synchrony, these findings on harmonicity lend support to the emerging view that parallel perceptual processes may govern auditory object formation in humans and non-human animals.

To our knowledge, this study is the first systematic investigation of harmonicity as an auditory object formation cue in frogs. There exists no single consensus on the behavioral significance of the prominent harmonic structures of the vocalizations used by many frogs. Using a physiological procedure called reflex modification, Simmons, (1988) demonstrated that males of *H. cinerea* exhibit sensitivity to shifts in the harmonic structure of two-tone complexes consisting of conspecific frequencies. However, behavioral investigations in *H. cinerea* using female phonotaxis by Gerhardt et al., (1990) and male evoked calling behavior by Simmons et al., (1993) demonstrated no significant differences in either phonotaxis or male evoked calling upon manipulating the harmonic relatedness of conspecific spectral components. One key difference between these two behavioral studies and the present study could be related to the test design. While Gerhardt et al., (1990) and Simmons et al., (1993) measured differential responsiveness to conspecific calls based on inharmonicity, the present study investigated perceptual segregation of conspecific and heterospecific signal components based on inharmonicity. Recall that in this study, conspecific spectral components were always harmonic with each other and instead their inharmonicity with the heterospecific spectral components was manipulated. Taking all of the above findings into consideration, we speculate that harmonicity may play a role in perceptual organization underlying species recognition but may not be behaviorally meaningful in the context of discriminating among conspecific mates or male-male competition with conspecifics.

### Multiple cue interactions during auditory object formation

Multiple cues may be differentially weighed, act in an additive fashion, or interact during auditory object formation. Most of our understanding of the effects of multiple cues on object formation comes from human psychoacoustic studies. Singh and Bregman (1997) showed that both common amplitude rise times and spectral similarity additively mediate integration of sounds. Elhilali et al. (2009) found that regardless of having considerable frequency separation (which promotes segregation), tones were integrated when they exhibited temporal synchrony. This finding indicated that temporal synchrony overrides spectral separation in mediating perceptual integration. Further, similar to the present study, Micheyl et al. (2013b) manipulated temporal coherence and harmonicity and found an additive effect of the two cues on perceptual organization. Recent findings from non-human animals suggest similar patterns for multiple cues mediating auditory object formation. Itatani and Klump (2020) showed that European starlings rely on the additive effects of common spatial origin and spectral similarity for perceptual organization, though spectral similarity had a larger effect compared to common spatial origin. Dent et al. (2016) showed, in zebra finches and budgerigars, stronger effects of spatial location and sound intensity than spectral cues in detecting a missing syllable from a stimulus bird song. Schwartz and Serratto Del Monte (2019) found no additional improvement in segregation when sounds that were spatially separated were also separated in their frequencies. The present study adds to this small but growing literature on auditory object formation in the presence of multiple cues in non-human animals. It is worth expanding the present findings by incorporating more auditory object formation cues. In future work, besides manipulating temporal onset asynchrony and inharmonicity, spatial location of the conspecific and heterospecific spectral components could also be manipulated using the present study design.

### Perceptual organization and neuroethological theories of auditory processing

An important finding to emphasize from this study was that sounds were integrated into the same auditory object *only if* they shared commonalities across multiple acoustic attributes, that is, both synchronous temporal onsets and harmonic relationships. It is worth discussing these findings in the light of how natural signals are structured and neuroethological theories of auditory processing. Natural signals often possess multiple spectro-temporal commonalities like harmonicity and slow temporal fluctuations that are correlated across frequency components (Kershenbaum et al., 2016; Nelken et al., 1999; Singh and Theunissen, 2003; Suga, 1992). These spectro-temporal commonalities are borne out of the physics of sound production. Neuroethological theories of auditory processing argue that neural systems in animals have evolved to process behaviorally relevant communication signals (Theunissen and Elie, 2014). These theories are supported by the discovery of neurons that respond best to features present in conspecific signals (Doupe and Konishi, 1991; Feng et al., 1990; Fitzpatrick et al., 1993; Grace et al., 2003; Römer, 2016). Consistent with this general view, auditory neurons in frogs are often tuned to sound frequencies emphasized in conspecific signals (reviewed in Feng et al., 1990; Simmons, 2013) and respond selectively to combinations of those frequencies (Fuzessery and Feng, 1983; Lee et al., 2017; Megela, 1983) or to species-specific temporal patterns (Edwards et al., 2002; Gupta et al., 2021).

Based on these neuroethological theories and the supporting evidence, auditory object formation may be based on the design principles of natural signals. That is, both temporal synchrony and harmonicity are required to promote perceptual integration because signal components in natural signals possess both of these regularities. This possibility could be tested by integrating various bioacoustic, behavioral and neurophysiological approaches in the same study system. For instance, the behavioral findings of the present study on *H. cinerea* could be compared against biologically realistic values of temporal synchrony and harmonicity present within and between signals of different males. Further, neurophysiological responses to the present experimental stimuli could be acquired to estimate the neural mechanisms of auditory object formation. Auditory processing at both the peripheral and central levels could be possible candidates for future neurophysiological investigations. Three distinct population of auditory neurons have been identified in the VIII nerve of *H. cinerea* that are tuned to low (< 500 Hz) mid (500-1200 Hz), and high (3100-3800 Hz) frequencies (Capranica et al., 1976; Moffat and Capranica, 1974). Ehret et al. (1983) found that responses of certain low frequency tuned VIII nerve fibers in *H. cinerea* exhibited a ‘two-tone’ suppression wherein certain inhibitory tones (comprising frequencies typical of heterospecific signals) suppressed the responses of those neurons to their excitatory frequencies. Further, neural recordings from the inferior colliculus and thalamus by Fuzessery and Feng, (1983) in *Rana pipiens*, Megela, (1983) in *Rana catesbeiana*, and Mudry and Capranica, (1987) and Lee et al., (2017) in *Hyla cinerea* showed the presence of certain combination-sensitive neurons that exhibited increased sensitivity at two distinct frequency regions and respond either maximally or exclusively to the combination of low and high frequencies present in the conspecific calls. Mudry and Capranica, (1987) demonstrated that combination sensitive neurons also exhibit suppression upon added stimulation by certain inhibitory frequencies (1200-1800 Hz; frequently reflected in the natural signals of heterospecific calls and also used as one of the heterospecific components in this study). This neural suppression of peripheral and central neurons in response to added stimulation by inhibitory frequencies is reflected in behavioral preferences too, as demonstrated in studies by Gerhardt, (1974b) and Gerhardt and Höbel, (2005). In those studies, adding a heterospecific spectral component to a conspecific call starkly reduced its attractiveness. We hypothesize that auditory object formation may be partially encoded by changes in the response patterns of the VIII nerve fibers and the combination sensitive neurons. Future studies could measure how shifts in temporal synchrony and harmonicity (comparable to the values used in this study) between the conspecific and heterospecific spectral components may introduce changes in the suppression patterns of the VIII nerve fibers and the combination sensitive neurons.

## Acknowledgements

We thank Isabelle Hoversten, Miranda Dahl and Madison Deile for their help collecting and testing frogs, and Jamie Hawkins from the Department of Natural Resources, Georgia and Scott Hamlin from Bowen Mills Fish Hatchery for generous access to collection sites. We are also grateful to Marlene Zuk, Andrew Oxenham and Trevor Wardill for feedback on this manuscript.

## Funding

This research was funded part by grants to MAB from the National Science Foundation (IOS-1452831 and IOS-2022253) and by grants and fellowships to LK from the University of Minnesota Graduate Program in Ecology, Evolution, and Behavior, the Bell Museum of Natural History, the University of Minnesota Graduate School and the Animal Behavior Society.

## Notes

### Competing Interest Statement

The authors have declared no competing interest.

## References

Akre, K. L. and Ryan, M. J. (2010). Proximity-dependent Response to Variably Complex Mating Signals in Túngara Frogs (Physalaemus pustulosus). Ethology 116, 1138–1145.

Bee, M. A. (2010). Spectral preferences and the role of spatial coherence in simultaneous integration in gray treefrogs (Hyla chrysoscelis). J. Comp. Psychol. 124, 412–424.

Bee, M. A. and Micheyl, C. (2008). The Cocktail Party Problem: What Is It? How Can It Be Solved? And Why Should Animal Behaviorists Study It? J. Comp. Psychol. 122, 235–251.

Bee, M. A. and Riemersma, K. K. (2008). Does common spatial origin promote the auditory grouping of temporally separated signal elements in grey treefrogs? Anim. Behav. 76, 831–843.

Bizley, J. K. and Cohen, Y. E. (2013). The what, where and how of auditory-object perception. Nat. Rev. Neurosci. 14, 693–707.

Bosch, J. and Márquez, R. (2002). Female preference function related to precedence effect in an amphibian anuran (Alytes cisternasii): tests with non-overlapping calls. Behav. Ecol. 13, 149–153.

Bregman, A. S. (1990). Auditory Scene Analysis: The Perceptual Organization of Sound.

Bregman, A. S. and Pinker, S. (1978). Auditory streaming and the building of timbre. Can. J. Psychol. Can. Psychol. 32, 19.

Broadbent, D. E. and Ladefoged, P. (1957). On the Fusion of Sounds Reaching Different Sense Organs. J. Acoust. Soc. Am. 29, 708–710.

Brokx, J. P. L. and Nooteboom, S. G. (1982). Intonation and the perceptual separation of simultaneous voices. J. Phon. 10, 23–36.

Brumm, H. and Slabbekoorn, H. (2005). Acoustic Communication in Noise. In Advances in the Study of Behavior, pp. 151–209. Academic Press.

Cai, H., Screven, L. A. and Dent, M. L. (2018). Behavioral measurements of auditory streaming and build-up by budgerigars ( Melopsittacus undulatus ) . J. Acoust. Soc. Am. 144, 1508–1516.

Capranica, R. R., Llinas, R. and Precht, W. (1976). Auditory system: morphology and physiology. Handb. Frog Neurobiol.

Dannenbring, G. L. and Bregman, A. S. (1978). Streaming vs. fusion of sinusoidal components of complex tones. Percept. Psychophys. 24, 369–376.

Darwin, C. J. (1981). Perceptual grouping of speech components differing in fundamental frequency and onset-time. Q. J. Exp. Psychol. Sect. A 33, 185–207.

Darwin, C. J. (1984). Perceiving vowels in the presence of another sound: Constraints on formant perception. J. Acoust. Soc. Am. 76, 1636–1647.

Darwin, C. J. and Ciocca, V. (1992). Grouping in pitch perception: Effects of onset asynchrony and ear of presentation of a mistuned component. J. Acoust. Soc. Am. 91, 3381–3390.

Darwin, C. J. and Gardner, R. B. (1986). Mistuning a harmonic of a vowel: Grouping and phase effects on vowel quality. J. Acoust. Soc. Am. 79, 838–845.

Darwin, C. J. and Gardner, R. B. (1987). Perceptual separation of speech from concurrent sounds. In The psychophysics of speech perception, pp. 112–124. Springer.

Darwin, C. J. and Hukin, R. W. (1998). Perceptual segregation of a harmonic from a vowel by interaural time difference in conjunction with mistuning and onset asynchrony. J. Acoust. Soc. Am. 103, 1080–1084.

Darwin, C. J. and Sutherland, N. S. (1984). Grouping frequency components of vowels: When is a harmonic not a harmonic? Q. J. Exp. Psychol. 36, 193–208.

Dent, M. L. and Bee, M. A. (2018). Principles of Auditory Object Formation by Nonhuman Animals.pp. 47–82. Springer, New York, NY.

Dent, M. L., Martin, A. K., Flaherty, M. M. and Neilans, E. G. (2016). Cues for auditory stream segregation of birdsong in budgerigars and zebra finches: Effects of location, timing, amplitude, and frequency. J. Acoust. Soc. Am. 139, 674–683.

Doupe, A. J. and Konishi, M. (1991). Song-selective auditory circuits in the vocal control system of the zebra finch. Proc. Natl. Acad. Sci. 88, 11339–11343.

Edwards, C. J., Alder, T. B. and Rose, G. J. (2002). Auditory midbrain neurons that count. Nat. Neurosci. 5, 934–936.

Ehret, G. and Gerhardt, H. C. (1980). Auditory masking and effects of noise on responses of the green treefrog (Hyla cinerea) to synthetic mating calls. J. Comp. Physiol. □ A 141, 13–18.

Ehret, G., Moffat, A. and Capranica, R. R. (1983). Two-tone suppression in auditory nerve fibers of the green treefrog (hyla cinerea). J. Acoust. Soc. Am. 73, 2093–2095.

Elhilali, M., Ma, L., Micheyl, C., Oxenham, A. J. and Shamma, S. A. (2009). Temporal Coherence in the Perceptual Organization and Cortical Representation of Auditory Scenes. Neuron 61, 317–329.

Farris, H. E., Rand, A. S. and Ryan, M. J. (2005). The effects of time, space and spectrum on auditory grouping in túngara frogs. J. Comp. Physiol. A. Neuroethol. Sens. Neural. Behav. Physiol. 191, 1173–1183.

Fay, R. R. (1998). Auditory stream segregation in goldfish (Carassius auratus). Hear. Res. 120, 69–76.

Feng, A. S., Hall, J. C. and Gooler, D. M. (1990). Neural basis of sound pattern recognition in anurans. Prog. Neurobiol. 34, 313–329.

Fenton, B., Jensen, F. H., Kalko, E. K. V. and Tyack, P. L. (2014). Sonar Signals of Bats and Toothed Whales. In (ed. Surlykke, A.), Nachtigall, P. E.), Fay, R. R.), and Popper, A. N.), pp. 11–59. New York, NY: Springer New York.

Fitzpatrick, D. C., Kanwal, J. S., Butman, J. A. and Suga, N. (1993). Combination-sensitive neurons in the primary auditory cortex of the mustached bat. J. Neurosci. 13, 931–940.

Fuzessery, Z. M. and Feng, A. S. (1983). Mating call selectivity in the thalamus and midbrain of the leopard frog (Rana p. pipiens): Single and multiunit analyses. J. Comp. Physiol. 150, 333–344.

Gardner, R. B., Gaskill, S. A. and Darwin, C. J. (1989). Perceptual grouping of formants with static and dynamic differences in fundamental frequency. J. Acoust. Soc. Am. 85, 1329–1337.

Gerhardt, H. C. (1974a). Behavioral Isolation of the Tree Frogs, Hyla cinerea and Hyla andersonii. Am. Midl. Nat. 91, 424.

Gerhardt, H. C. (1974b). The significance of some spectral features in mating call recognition in the green treefrog (Hyla cinerea). J. Exp. Biol. 61, 229–241.

Gerhardt, H. C. (1976). Significance of two frequency bands in long distance vocal communication in the green treefrog. Nature 261, 692–694.

Gerhardt, H. C. (1981). Mating call recognition in the green treefrog (Hyla cinerea): Importance of two frequency bands as a function of sound pressure level. J. Comp. Physiol. □ A 144, 9–16.

Gerhardt, H. C. (1987). Evolutionary and neurobiological implications of selective phonotaxis in the green treefrog, Hyla cinerea. Anim. Behav. 35, 1479–1489.

Gerhardt, H. C. (2001). Acoustic communication in two groups of closely related treefrogs. In Advances in the Study of Behavior, pp. 99–167. Academic Press.

Gerhardt, H. C. and Höbel, G. (2005). Mid-frequency suppression in the green treefrog (Hyla cinerea): Mechanisms and implications for the evolution of acoustic communication. *J. Comp. Physiol. A Neuroethol. Sensory, Neural*, Behav. Physiol. 191, 707–714.

Gerhardt, H. C. and Huber, F. (2002). Acoustic communication in insects and anurans : common problems and diverse solutions. Chicago: University of Chicago Press.

Gerhardt, H. C., Allan, S. and Schwartz, J. J. (1990). Female green treefrogs (Hyla cinerea) do not selectively respond to signals with a harmonic structure in noise. J. Comp. Physiol. A 166, 791–794.

Grace, J. A., Amin, N., Singh, N. C. and Theunissen, F. E. (2003). Selectivity for conspecific song in the zebra finch auditory forebrain. J. Neurophysiol. 89, 472–487.

Grafe, T. U. (1999). A function of synchronous chorusing and a novel female preference shift in an anuran. Proc. R. Soc. London. Ser. B Biol. Sci. 266, 2331–2336.

Greenfield, M. D. (1994). Cooperation and conflict in the evolution of signal interactions. Annu. Rev. Ecol. Syst. 25, 97–126.

Greenfield, M. D. (2005). Mechanisms and Evolution of Communal Sexual Displays in Arthropods and Anurans. Adv. Study Behav. 35, 1–62.

Greenfield, M. D., Tourtellot, M. K. and Snedden, W. A. (1997). Precedence effects and the evolution of chorusing. Proc. R. Soc. London. Ser. B Biol. Sci. 264, 1355–1361.

Griffiths, T. D. and Warren, J. D. (2004). What is an auditory object? Nat. Rev. Neurosci. 5, 887–892.

Gupta, S. and Bee, M. A. (2020). Treefrogs exploit temporal coherence to form perceptual objects of communication signals. Biol. Lett. 16, 20200573.

Gupta, S., Alluri, R. K., Rose, G. J. and Bee, M. A. (2021). Neural basis of acoustic species recognition in a cryptic species complex. J. Exp. Biol. 224, jeb243405.

Hall, M. L. (2009). A review of vocal duetting in birds. Adv. Study Behav. 40, 67–121.

Hardin, J. W. and Hilbe, J. M. (2012). Generalized estimating equations. 2nd edn Boca Raton. FL Chapman Hall/CRC.

Hartmann, W. M., McAdams, S. and Smith, B. K. (1990). Hearing a mistuned harmonic in an otherwise periodic complex tone. J. Acoust. Soc. Am. 88, 1712–1724.

Höbel, G. (2015). Sexual differences in responses to cross-species call interference in the green treefrog (Hyla cinerea). Behav. Ecol. Sociobiol. 69, 695–705.

Höbel, G. and Gerhardt, H. C. (2003). Reproductive character displacement in the acoustic communication system of green tree frogs (Hyla cinerea). Evolution (N. Y). 57, 894–904.

Höbel, G. and Gerhardt, H. C. (2007). Sources of selection on signal timing in a tree frog. Ethology 113, 973–982.

Hoeschele, M. (2017). Animal pitch perception: Melodies and harmonies. Comp. Cogn. Behav. Rev. 12, 5.

Hua, X., Fu, C., Li, J., de Oca, A. N. M. and Wiens, J. J. (2009). A revised phylogeny of Holarctic treefrogs (genus Hyla) based on nuclear and mitochondrial DNA sequences. Herpetologica 65, 246–259.

Hukin, R. W. and Darwin, C. J. (1995). Comparison of the effect of onset asynchrony on auditory grouping in pitch matching and vowel identification. Percept. Psychophys. 57, 191–196.

Hulse, S. H. (2002). Auditory scene analysis in animal communication. Adv. Study Behav. 31, 163–200.

Hyland Bruno, J. and Tchernichovski, O. (2019). Regularities in zebra finch song beyond the repeated motif. Behav. Processes 163, 53–59.

Itatani, N. and Klump, G. M. (2020). Interaction of spatial and non-spatial cues in auditory stream segregation in the European starling. Eur. J. Neurosci. 51, 1191–1200.

Jones, D. L., Jones, R. L. and Ratnam, R. (2014). Calling dynamics and call synchronization in a local group of unison bout callers. *J. Comp. Physiol. A Neuroethol. Sensory, Neural*, Behav. Physiol. 200, 93–107.

Kalra, L., Altman, S. and Bee, M. A. (2024). Perceptually salient differences in a species recognition cue do not promote auditory streaming in eastern grey treefrogs (Hyla versicolor). J. Comp. Physiol. A.

Kershenbaum, A., Blumstein, D. T., Roch, M. A., Akçay, Ç., Backus, G., Bee, M. A., Bohn, K., Cao, Y., Carter, G., Cäsar, C., et al. (2016). Acoustic sequences in non-human animals: a tutorial review and prospectus. Biol. Rev. Camb. Philos. Soc. 91, 13–52.

Klinge, A. and Klump, G. M. (2009). Frequency difference limens of pure tones and harmonics within complex stimuli in Mongolian gerbils and humans. J. Acoust. Soc. Am. 125, 304–314.

Klinge, A. and Klump, G. (2010). Mistuning detection and onset asynchrony in harmonic complexes in Mongolian gerbils. J. Acoust. Soc. Am. 128, 280–290.

Klump, G. M. and Gerhardt, H. C. (1992). Mechanisms and Function of Call-Timing in Male-Male Interactions in Frogs. In Playback and Studies of Animal Communication (ed. McGregor, P. K.), pp. 153–174. Boston, MA: Springer US.

Ladefoged, P. and Disner, S. F. (2012). Vowels and consonants. John Wiley & Sons.

Lee, N., Schrode, K. M. and Bee, M. A. (2017). Nonlinear processing of a multicomponent communication signal by combination-sensitive neurons in the anuran inferior colliculus. J. Comp. Physiol. A 203, 749–772.

Lee, N., Christensen-Dalsgaard, J., White, L. A., Schrode, K. M. and Bee, M. A. (2021). Lung mediated auditory contrast enhancement improves the signal-to-noise ratio for communication in frogs. Curr. Biol. 31, 1488–1498.

Legett, H. D., Aihara, I. and Bernal, X. E. (2020). Signal synchrony and alternation among neighbor males in a Japanese stream breeding treefrog, Buergeria japonica. Curr. Herpetol. 39, 80–85.

Legett, H. D., Aihara, I. and Bernal, X. E. (2021). The dual benefits of synchronized mating signals in a Japanese treefrog: attracting mates and manipulating predators. Philos. Trans. R. Soc. B 376, 20200340.

Lohr, B. and Dooling, R. J. (1998). Detection of changes in timbre and harmonicity in complex sounds by zebra finches (Taeniopygia guttata) and budgerigars (Melopsittacus undulatus). J. Comp. Psychol. 112, 36.

Ma, L., Micheyl, C., Yin, P., Oxenham, A. J. and Shamma, S. A. (2010). Behavioral measures of auditory streaming in ferrets (Mustela putorius). J. Comp. Psychol. 124, 317–330.

Madsen, S. M. K. and Moore, B. C. J. (2014). Effects of compression and onset/offset asynchronies on the detection of one tone in the presence of anothera). J. Acoust. Soc. Am. 135, 2902–2912.

McDermott, J. H. (2009). The cocktail party problem. Curr. Biol. 19, R1024–R1027.

McDermott, J. H. and Oxenham, A. J. (2008). Music perception, pitch, and the auditory system. Curr. Opin. Neurobiol. 18, 452–463.

Mecham, J. S. (1965). Genetic relationships and reproductive isolation in southeastern frogs of the genera Pseudacris and Hyla. Am. Midl. Nat. 269–308.

Megela, A. L. (1983). Auditory response properties of the anuran thalamus: nonlinear facilitation. In Advances in vertebrate neuroethology, pp. 895–899. Springer.

Micheyl, C., Hanson, C., Demany, L., Shamma, S. and Oxenham, A. J. (2013a). Auditory stream segregation for alternating and synchronous tones. J. Exp. Psychol. Hum. Percept. Perform. 39, 1568.

Micheyl, C., Kreft, H., Shamma, S. and Oxenham, A. J. (2013b). Temporal coherence versus harmonicity in auditory stream formation. J. Acoust. Soc. Am. 133, EL188–EL194.

Minckley, R. L. and Greenfield, M. D. (1995). Psychoacoustics of female phonotaxis and the evolution of male signal interactions in Orthoptera. Ethol. Ecol. Evol. 7, 235–243.

Moffat, A. J. M. and Capranica, R. R. (1974). Sensory Processing in the Peripheral Auditory System of Treefrogs (Hyla). J. Acoust. Soc. Am. 55, 480.

Moore, B. C. J. and Gockel, H. E. (2012). Properties of auditory stream formation. Philos. Trans. R. Soc. B Biol. Sci. 367, 919–931.

Moore, B. C. J., Peters, R. W. and Glasberg, B. R. (1985). Thresholds for the detection of inharmonicity in complex tones. J. Acoust. Soc. Am. 77, 1861–1867.

Moore, B. C. J., Glasberg, B. R. and Peters, R. W. (1986). Thresholds for hearing mistuned partials as separate tones in harmonic complexes. J. Acoust. Soc. Am. 80, 479–483.

Mudry, K. M. and Capranica, R. R. (1987a). Correlation between auditory evoked responses in the thalamus and species-specific call characteristics: I. Rana catesbeiana (Anura: Ranidae). J. Comp. Physiol. A 160, 477–489.

Mudry, K. M. and Capranica, R. R. (1987b). Correlation between auditory thalamic area evoked responses and species-specific call characteristics. II. H. Hyla cinerea (Anura: Hylidae). J. Comp. Physiol. A. 161, 407–416.

Neilans, E. G. and Dent, M. L. (2015). Temporal coherence for complex signals in budgerigars (Melopsittacus undulatus) and humans (Homo sapiens). J. Comp. Psychol. 129, 174.

Nelken, I., Rotman, Y. and Yosef, O. B. (1999). Responses of auditory-cortex neurons to structural features of natural sounds. Nature 397, 154–157.

Nityananda, V. and Bee, M. A. (2011). Finding your mate at a cocktail party: Frequency separation promotes auditory stream segregation of concurrent voices in multi-species frog choruses. PLoS One 6,.

Oldham, R. S. and Gerhardt, H. C. (1975). Behavioral Isolating Mechanisms of the Treefrogs Hyla cinerea and H. gratiosa. Copeia 1975, 223.

Prestwich, K. N. (1994). The energetics of acoustic signaling in anurans and insects. Integr. Comp. Biol. 34, 625–643.

Reichert, M. S., Luttbeg, B. and Hobson, E. A. (2024). Collective signalling is shaped by feedbacks between signaller variation, receiver perception and acoustic environment in a simulated communication network. Philos. Trans. B 379, 20230186.

Roberts, B. and Moore, B. C. J. (1991). The influence of extraneous sounds on the perceptual estimation of first-formant frequency in vowels under conditions of asynchrony. J. Acoust. Soc. Am. 89, 2922–2932.

Robillard, T., Höbel, G. and Carl Gerhardt, H. (2006). Evolution of advertisement signals in North American hylid frogs: vocalizations as end-products of calling behavior. Cladistics 22, 533–545.

Römer, H. (2016). Matched filters in insect audition: tuning curves and beyond. Ecol. Anim. Senses Matched Filters Econ. Sens. 83–109.

Schul, J. and Sheridan, R. A. (2006). Auditory stream segregation in an insect. Neuroscience 138, 1–4.

Schwartz, J. J. (1987). The function of call alternation in anuran amphibians: a test of three hypotheses. Evolution (N. Y). 41, 461–471.

Schwartz, J. J. and Gerhardt, H. C. (1989). Spatially mediated release from auditory masking in an anuran amphibian. J. Comp. Physiol. A 166, 37–41.

Schwartz, J. J. and Gerhardt, H. C. (1995). Directionality of the auditory system and call pattern recognition during acoustic interference in the gray treefrog, Hyla versicolor. Audit. Neurosci. 1, 195–206.

Schwartz, J. J. and Serratto Del Monte, M. E. (2019). Spatially-mediated call pattern recognition and the cocktail party problem in treefrog choruses: can call frequency differences help during signal overlap? Bioacoustics 28, 312–328.

Simmons, A. M. (1988). Selectivity for harmonic structure in complex sounds by the green treefrog (Hyla cinerea). J. Comp. Physiol. A 162, 397–403.

Simmons, A. M. (2013). “To Ear is Human, to Frogive is Divine”: Bob Capranica’s legacy to auditory neuroethology. J. Comp. Physiol. A 199, 169–182.

Simmons, A. M. and Bean, M. E. (2000). Perception of mistuned harmonics in complex sounds by the bullfrog (Rana catesbeiana). J. Comp. Psychol. 114, 167.

Simmons, A. M., Buxbaum, R. C. and Mirin, M. P. (1993). Perception of complex sounds by the green treefrog, Hyla cinerea: envelope and fine-structure cues. J. Comp. Physiol. A 173, 321–327.

Singh, P. G. and Bregman, A. S. (1997). The influence of different timbre attributes on the perceptual segregation of complex-tone sequences. J. Acoust. Soc. Am. 102, 1943–1952.

Singh, N. C. and Theunissen, F. E. (2003). Modulation spectra of natural sounds and ethological theories of auditory processing. J. Acoust. Soc. Am. 114, 3394–3411.

Snedden, W. A. and Greenfield, M. (1998). Females prefer leading males: relative call timing and sexual selection in katydid choruses. Anim. Behav. 56, 1091–1098.

Suga, N. (1992). Philosophy and Stimulus Design for Neuroethology of Complex-Sound Processing. Philos. Trans. R. Soc. London. Ser. B. Biol. Sci. 336, 423–428.

Theunissen, F. E. and Elie, J. E. (2014). Neural processing of natural sounds. Nat. Rev. Neurosci. 15, 355–366.

Thorpe, W. H. (1975). The biological significance of duetting and antiphonal song. Acta Neurobiol. Exp. (Wars). 35, 517–528.

Tuttle, M. D. and Ryan, M. J. (1982). The role of synchronized calling, ambient light, and ambient noise, in anti-bat-predator behavior of a treefrog. Behav. Ecol. Sociobiol. 11, 125–131.

Wells, K. D. and Schwartz, J. J. (1984). Vocal communication in a neotropical treefrog, Hyla ebraccata: advertisement calls. Anim. Behav. 32, 405–420.

Winn, H. E., Thompson, T. J., Cummings, W. C., Hain, J., Hudnall, J., Hays, H. and Steiner, W. W. (1981). Song of the humpback whale - Population comparisons. Behav. Ecol. Sociobiol. 8, 41–46.

