## Supplementary text for "Treefrogs exploit temporal onset synchrony and harmonicity in forming auditory objects of vocal communication signals"

**Supplementary materials**

**Fig. S1** Proportion of subjects choosing the standard call across all combinations of temporal onset asynchrony (ΔT) compared between ΔF = +10% and ΔF = -10%. The dashed horizontal line indicates the null expectation of 0.50 in a two-alternative choice tests. Gray and black bars indicate ΔF = +10% and ΔF = -10%, respectively. Error bars represent 95% exact binomial confidence intervals.

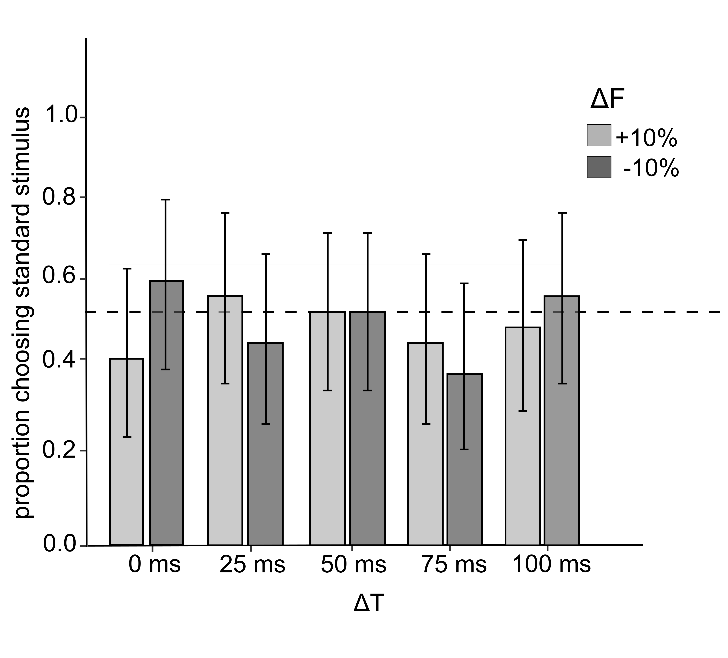

| **Supplementary Table 1:** Results of the Generalized Estimating Equation (GEE) model. Here, only the data for which ΔF = ±10% was analyzed to compare ΔF = +10% and ΔF = -10%. The effects of temporal onset asynchrony (ΔT), inharmonicity (ΔF), and their interaction are reported in relation to the reference test where ΔT = 0 ms and ΔF = -10%. Condition represents any manipulation in ΔT, ΔF or the interactive effects of different combinations of ΔT and ΔF. β indicates the estimate value for each condition. Wald Χ^2^ and P values correspond to the results of the Wald-statistics. | | | |
| --- | --- | --- | --- |
| **Condition** | **β** | **Wald Χ^2^** | ***P* value** |
| ΔT = 25 ms | -0.54 | 1.01 | 0.32 |
| ΔT = 50 ms | -0.27 | 0.25 | 0.62 |
| ΔT = 75 ms | -0.82 | 3.15 | 0.08 |
| ΔT = 100 ms | -0.14 | 0.09 | 0.76 |
| ΔF = +10% | -0.67 | 1.65 | 0.20 |
| ΔT = 25 ms × ΔF = +10% | 1.10 | 2.16 | 0.14 |
| ΔT = 50 ms × ΔF = +10% | 0.67 | 0.94 | 0.33 |
| ΔT = 75 ms × ΔF = +10% | 0.95 | 1.99 | 0.16 |
| ΔT = 100 ms × ΔF = +10% | 0.41 | 0.36 | 0.55 |
